# A high-affinity RBD-targeting nanobody improves fusion partner’s potency against SARS-CoV-2

**DOI:** 10.1101/2020.09.24.312595

**Authors:** Hebang Yao, Hongmin Cai, Tingting Li, Bingjie Zhou, Wenming Qin, Dimitri Lavillette, Dianfan Li

## Abstract

A key step to the SARS-CoV-2 infection is the attachment of its Spike receptor-binding domain (S RBD) to the host receptor ACE2. Considerable research have been devoted to the development of neutralizing antibodies, including llama-derived single-chain nanobodies, to target the receptor-binding motif (RBM) and to block ACE2-RBD binding. Simple and effective strategies to increase potency are desirable for such studies when antibodies are only modestly effective. Here, we identify and characterize a high-affinity synthetic nanobody (sybody, SR31) as a fusion partner to improve the potency of RBM-antibodies. Crystallographic studies reveal that SR31 binds to RBD at a conserved and ‘greasy’ site distal to RBM. Although SR31 distorts RBD at the interface, it does not perturb the RBM conformation, hence displaying no neutralizing activities itself. However, fusing SR31 to two modestly neutralizing sybodies dramatically increases their affinity for RBD and neutralization activity against SARS-CoV-2 pseudovirus. Our work presents a tool protein and an efficient strategy to improve nanobody potency.

## Introduction

SARS-CoV-2, the pathogenic virus for COVID-19, has caused a global pandemic since its first report in early December 2019 in Wuhan China (*1*), posing a gravely crisis for health and economic and social order. SARS-CoV-2 is heavily decorated by its surface Spike (S) (*2, 3*), a single-pass membrane protein that is key for the host-virus interactions. During the infection, S is cleaved by host proteases (*4, 5*), yielding the N-terminal S1 and the C-terminal S2 subunit. S1 binds to angiotensin-converting enzyme 2 (ACE2) (*6-10*) on the host cell membrane via its receptor-binding domain (RBD), causing conformational changes that trigger a secondary cleavage needed for the S2-mediated membrane fusion at the plasma membrane or in the endosome. Because of this essential role, RBD has been a hot spot for the development of therapeutic monoclonal antibodies (mAbs) and vaccine (*11-28*).

Llama-derived heavy chain-only antibodies (nanobodies) are attractive bio-therapeutics (*29*). These small (∼14 kDa) proteins are robust, straightforward to produce, and amenable to engineering such as mutation and fusion. Owing to their ultra-stability, nanobodies have been reported to survive nebulization, a feature that has been explored for the development of inhaled nanobodies to treat respiratory viral diseases (*30, 31*) which categorizes COVID-19. Owing to their high sequence similarities with human type 3 VH domains (VH3), nanobodies are known to cause little immunogenicity (*29*). For the same reason, they can be humanized with relative ease to reduce immunogenicity when needed. Therefore, nanobodies as biotherapeutics are being increasingly recognized. Examples of nanobody drugs include caplacizumab (*32*) for the treatment of acquired thrombotic thrombocytopenic purpura, and ozoralizumab and vobarilizumab that are in the clinical trials for rheumatoid arthritis (*29, 33*). Recently, several groups have independently reported neutralizing nanobodies (*22, 34-39*) or single-chain VH antibodies (*40*) against SARS-CoV-2 with variable potencies.

We have recently reported several synthetic nanobodies (sybodies) which bind RBD with various affinity and neutralizing activity (*35*). Affinity and neutralizing activity are very important characteristics for therapeutic antibodies, and they can be improved by a number of ways such as random mutagenesis (*22, 36*) and structure-based design. Previously, in the case of one modestly-neutralizing sybody MR17, we have determined its structure and designed a single mutant that improves its potency by over 23 folds (*35*). The rational design approach, while very effective, inevitably requires high-resolution structural information which are non-trivial to obtain. Generally applicable tools will be welcome.

Here, we report a strategy to increase sybody potency by biparatopic fusion with SR31, a sybody that binds RBD tightly with a *K*_D_ of 5.6 nM. As revealed by crystal structure, SR31 engages the RBD at a conserved site that is distal to the RBM. As such, it does not neutralize SARS-CoV-2 but forms non-competing pairs with several other RBM-binders and increases their neutralization potency when conjugated. SR31 may be used as a general affinity-enhancer for both detection and therapeutic applications.

## Results and Discussion

### A high-affinity RBD binder without neutralizing activity

Previously, we generated 99 sybodies from three highly diverse synthetic libraries by ribosome and phage display with *in vitro* selections against the SARS-CoV-2 RBD. Most of the sybody binders showed neutralizing activity. Interestingly, about 10 sybodies bind RBD but showed no neutralizing activities (*35*) even at 1 μM concentration.

One such sybodies, named SR31, was characterized in this study. In analytic fluorescence-detection size exclusion chromatography (FSEC), SR31 caused earlier retention of RBD (**Fig. 1A**) which was included at a low concentration (0.5 μM), suggesting nanomolar affinity for SR31-RBD binding. This was confirmed by bio-layer interferometry analysis (**Fig. 1B**) which showed a *K*_D_ of 5.6 nM and an off-rate of 1 × 10^−3^ s^-1^. Consistent with its inability to neutralize SARS-CoV-2 pseudovirus, SR31 did not affect RBD-ACE2 binding (**Fig. 1C**).

**Fig. 1.**
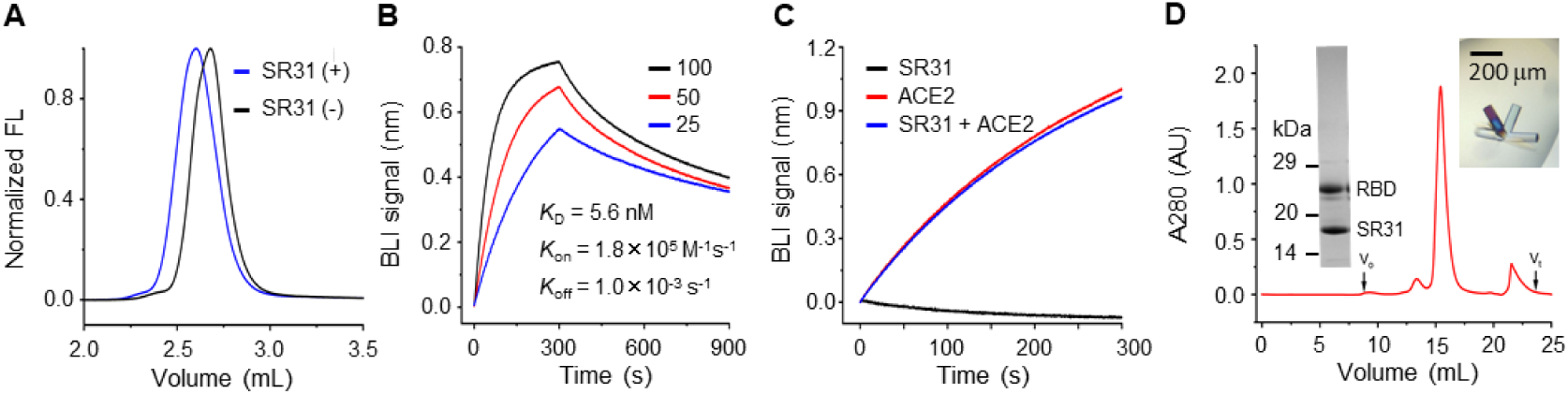
Characterization of the SR31-RBD complex. (**A**) FSEC of RBD in the absence (black) and presence (blue) of SR31. (**B**) Bio-layer interferometry (BLI) assay with RBD immobilized and SR31 as analyte at three concentrations (nM). (**C**) SR31 does not inhibit ACE2 binding. A RBD-coated sensor saturated with SR31 was soaked in 50 nM of SR31 with (blue) and without (black) 25 nM ACE2. As a control, the assay was performed with RBD immobilized and ACE2 as analyte (red). (**D**) Purification (SEC and SDS-PAGE) and crystallization of the RBD-SR31 complex.

### Structure of SR31 in complex with RBD

To characterize the SR31-RBD interactions in detail, we purified the complex (**Fig. 1D**), and obtained crystals (**Fig. 1D**) that diffracted to 1.97 Å resolution (**Table 1**). The structure was solved by molecular replacement using the published RBD and sybody structures (PDB IDs 6M0J and 5M13) (*6, 41*) as search models. The structure was refined to *R*_work_/*R*_free_ of 0.182/0.207 (**Table 1**). The asymmetric unit contained one molecule each for the RBD and SR31, indicating an expected 1:1 stoichiometry.

**Table 1.** Data collection and refinement statistics.

|  | SR31 + RBD | SR31-MR17 + RBD |
| --- | --- | --- |
| **Data collection** |  |  |
| Space group | P 3<sub>1</sub> 2 1 | P 6<sub>5</sub> 2 2 |
| Cell dimensions |  |  |
| *a*, *b*, *c* (Å) | 92.39, 92.39, 101.15 | 73.38, 73.38, 478.36 |
| *α*, *β*, *γ* (°) | 90, 90, 120 | 90, 90, 120 |
| Wavelength (Å) | 0.97854 | 0.98754 |
| Resolution (Å) | 19.61 – 1.97<br>(2.04 – 1.97)<sup>a</sup> | 49.70- 2.10 (2.16-2.10) |
| *R*<sub>merge</sub> | 0.091 (1.425) | 0.140 (1.409) |
| *R*<sub>pim</sub> | 0.209 (0.336) | 0.034 (0.373) |
| *I*/σ*I* | 19.5 (1.7) | 12.6 (2.0) |
| Completeness (%) | 99.9 (100.0) | 99.7 (96.6) |
| Multiplicity | 19.8 (18.8) | 18.6 (14.5) |
| *CC* * <sup>b</sup> | 1.000 (0.949) | 0.999 (0.965) |
| **Refinement** |  |  |
| Resolution (Å) | 19.61 – 1.97 | 49.70 - 2.10 |
| No. reflections | 35,702 | 46,078 |
| *R*<sub>work</sub> / *R*<sub>free</sub> | 0.1822 / 0.2071 | 0.1949 / 0.2359 |
| No. atoms | 2,916 | 3,892 |
| Protein | 2,592 | 3,437 |
| Ligands | 158 | 235 |
| Solvent | 166 | 220 |
| No. residues | 329 | 435 |
| B-factors (Å<sup>2</sup>) | 49.49 | 50.52 |
| Protein | 48.01 | 48.17 |
| Ligand/ion | 73.11 | 78.91 |
| Solvent | 50.19 | 56.78 |
| R.m.s deviations |  |  |
| Bond lengths (Å) | 0.008 | 0.008 |
| Bond angles (°) | 0.870 | 0.830 |
| Ramachandran |  |  |
| Favoured (%) | 96.62 | 97.18 |
| Allowed (%) | 3.38 | 2.82 |
| Outlier (%) | 0 | 0 |
| **PDB ID** | 7D2Z | 7D30 |

<sup>a</sup>Highest resolution shell is shown in parenthesis. <sup>b</sup> $CC^* = \sqrt{\frac{2CC_{1/2}}{1+CC_{1/2}}}$

SR31 binds to the RBD sideways at a buried surface area of 1,386.3 Å^2^ (**Fig. 2A**), which is significantly larger than that for the previously reported sybodies SR4 (727.4 Å^2^) and MR17 (853.944 Å^2^) (*35*). The binding surface is near a heavily decorated glycosylation site, Asn343 (**Fig. 2A-2C**), which, although at an apparent strategic position to possibly divide the accessible surfaces for immune surveillance, does not show clashes with SR31. All three CDRs participated in the interaction by providing five (CDR1), three (CDR2), and nine H-bonds (CDR3) (**Fig. 2E-2G**). Peculiarly, the CDR3, which contains a cluster of hydrophobic side chains that include Met99, Val100, Phe102, Trp103, and Tyr104, inserted into a greasy pocket (**Fig. 2B**) in the RBD that was lined with twelve hydrophobic/aromatic residues (**Fig. 2F**). Unlike salt bridges, hydrophobic interactions are more tolerant to environment such as change of pH and ionic strength. In addition, they are less specific and thus less likely to be affected by mutations. This binding mode thus makes SR31 an attractive candidate for detection purposes.

**Fig. 2.**
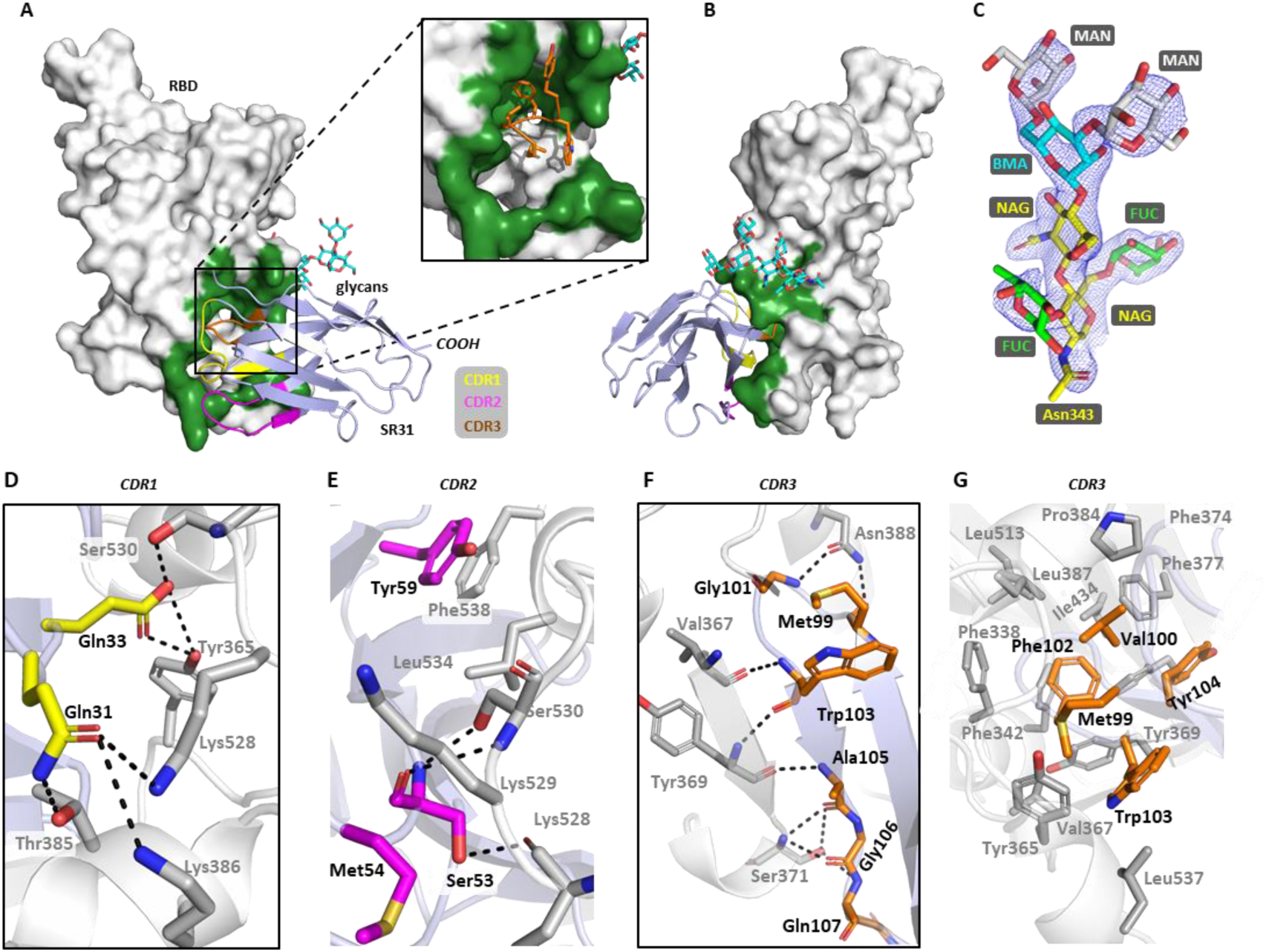
Crystal structure of the SR31-RBD complex. (**A**) The overall structure of SR31 (light blue) in complex with RBD (grey) which contains Asn343-linked glycans (cyan). The expanded view highlights a deep hydrophobic pocket for CDR3 binding. (**B**) The overall structure viewed at a different angle. (**C**) 2*Fo-Fc* map of the Asn343-linked glycans. MAN, mannose; BMA, β-_D_-mannose; FUC, fucose; NAG, N-acetylglucosamine. (**D-G**) Detailed interactions between RBD and the CDR1 (**D**), CDR2 (**E**), and CDR3 (**F, G**). The hydrophobic network formed between CDR3 (orange) and the hydrophobic pocket in RBD (grey) is shown in **G**. Residues from SR31 are labeled with black texts and residues from RBD are labeled with grey texts. Dash lines indicate hydrogen bonds or salt bridges within 3.6 Å.

Most RBD-targeting neutralizing antibodies, including neutralizing nanobodies characterized so far (*8, 13-15, 19, 20, 22-24, 26-28, 34, 35, 37*), engage the RBD at the receptor-binding motif (RBM) (**Fig. 3A**), thus competing off ACE2 and preventing viral entry. Aligning the ACE2 structure to the SR31-RBD structure showed that the SR31-binding epitope is distant from the RBM (**Fig. 3A**). Comparing the epitopes of existing monoclonal antibodies showed that the SR31 epitope partly overlaps with CR3022 (*12*) and the recently identified EY6A (*22*) (**Fig. 3B, 3C**). It has been established that the binding of the bulky CR3022 and EY6A at the interface between RBD and the N-terminal domain (NTD) of the adjacent monomer destabilizes the S trimer and converts the pre-fusion conformation to the infection-incompetent post-fusion state, thus conferring neutralization activity (*21, 22*). Despite the epitope overlapping, SR31 approaches RBD at a different angle to that of CR3022 (**Fig. 3C**). This angular difference, together with its minute size, may allow SR31 to bind two of the three sites in the ‘open’-S (*3*): the ‘up-RBD’, and the ‘down-RBD’ at the clockwise monomer (**Fig. 3D**). Taken together, the structural data rationalize the high-affinity binding between SR31 and RBD, and its inability to neutralize SARS-CoV-2.

**Fig. 3.**
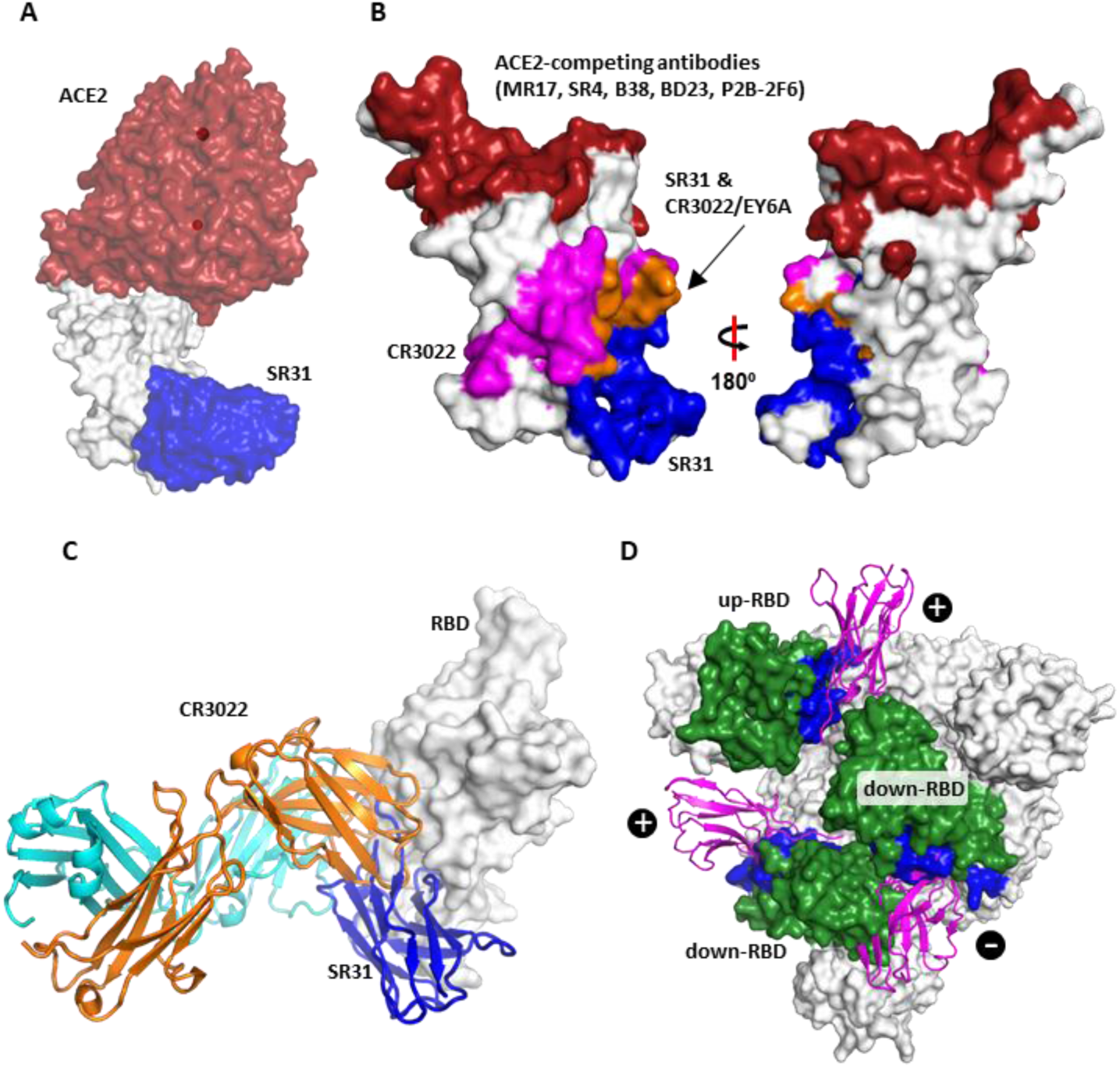
The SR31 epitope. (**A**) SR31 (blue) binds to RBD (grey) at a surface distal to the binding site of ACE2 (red). (**B**) Comparison of the SR31 epitope with epitopes for other RBD-targeting nanobodies *(22, 35, 36, 39)* and mAbs (*13-15, 19, 20, 23, 24, 26-28*). Red, the collective epitope of RBM-binders; blue, the SR31 epitope; magenta, the collective epitope of CR3022 and EY6A; orange, the overlap between the CR3022/EY6A and SR31 epitope. (**C**) SR31 (blue) binds RBD (grey) at a surface that overlaps with the epitope of CR3022 (orange and cyan) but approaches RBD at a different angle. (**D**) The binding site of SR31 in the context of the S trimer at its pre-fusion ‘open’ state with one RBD in the ‘up’ conformation and two in the ‘down’ conformation. The structure (PDB ID 6yvb) (*3*) is viewed from the ‘top’ (perpendicular to the viral membrane). The SR31 epitope is shown in blue. The three RBDs are colored green. SR31 (magenta cartoon) was aligned to the S trimer (surface presentation) by superposing the SR31-RBD structure to each of the RBD. ‘+’, no or minor clashes; ‘-’, with severe clashes.

Because nanobodies are relatively easy to produce, the availability of nanobodies that recognize a wide spectrum of epitopes can be a useful toolkit to probe binding mode of uncharacterized antibodies using competitive binding assays. They may also be used to select binders with new epitopes by including them as pre-formed sybody-RBD complexes during *in vitro* selection (and thus excluding binders at the same site).

### SR31-RBD structure suggests high RBD domain stability

Structure alignment of SR31-RBD with ACE2-RBD revealed that the two RBD structures were overall very similar with a Cα RMSD of 0.452 Å (**Fig. 4A**). Nevertheless, significant structural rearrangements at the binding interface were observed (**Fig. 4A, 4B**). Specifically, the small α-helix α_364-370_ (numbers mark start-end) moves towards the direction of RBM by a dramatic ∼8.0 Å and transforms to a short β-sheet (β_367-370_) which in turn forms a parallel β-sheet pair with β_102-104_ in the CDR3 region. In addition, nudged by the CDR1, the short helix α_383-388_ swings towards the RBD core by ∼4.0 Å.

**Fig. 4.**
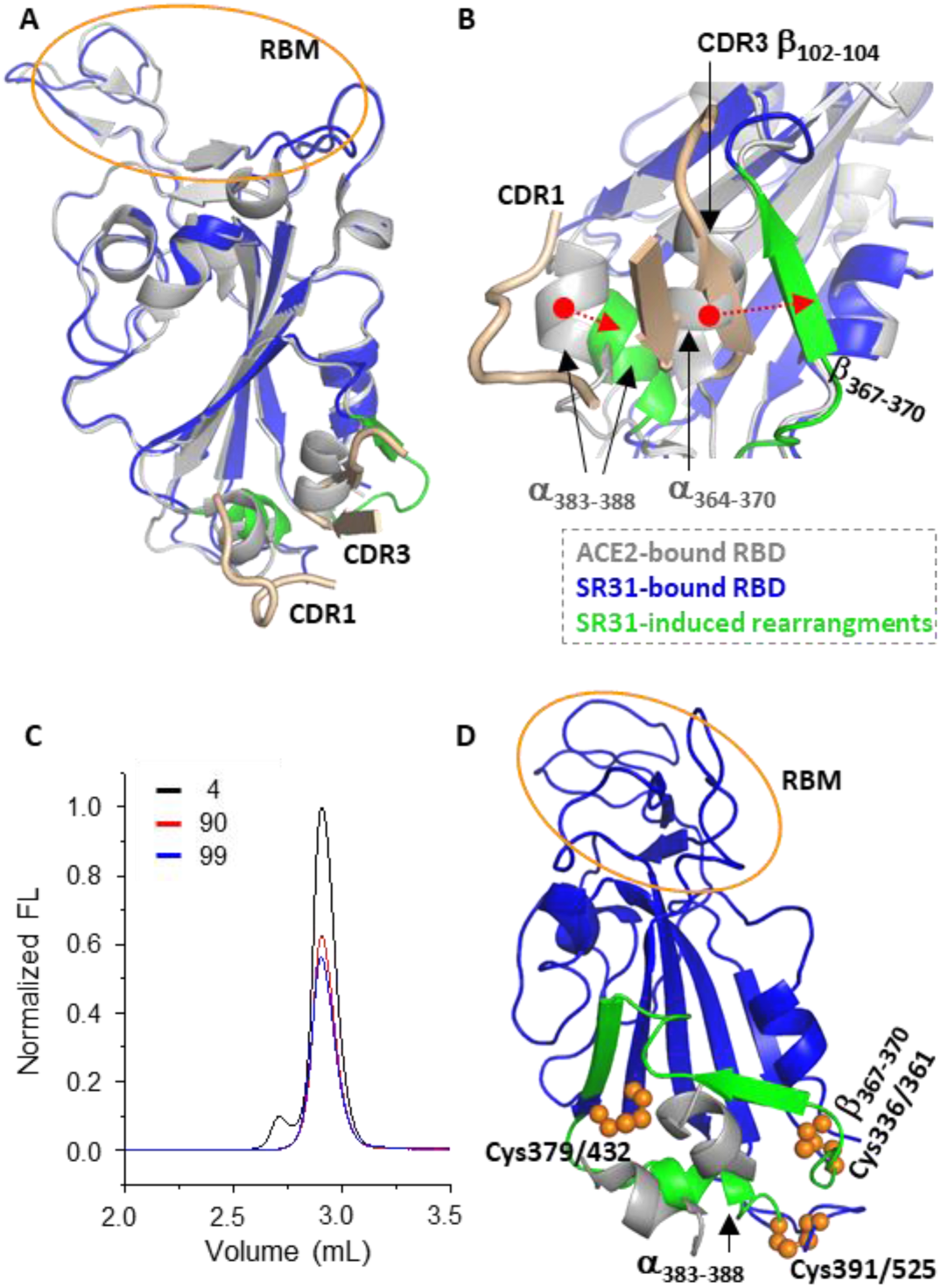
RBD is a very stable domain. (**A, B**) The overview (**A**) and expanded view (**B**) of the comparison between the ACE2-bound RBD (grey) and SR31-bound RBD (blue). SR31-binding deforms the RBD at the binding site (green) but not at the RBM region (yellow circle). The two SR31 CDRs involved in the deformation are shown as wheat cartoon. In **B**, two structure rearrangements (green) are shown at a different angle. The α_383-388_ helix in the ACE2-bound form is pushed towards the RBD core; and the short helix α_364-370_ is transformed into a β-strand (β_367-370_) which forms a parallel β-sheet with β_102-104_ from SR31 CDR3. (**C**) An indirect stability assay of the RBD using fluorescence-detection size exclusion chromatography. The RBD was incubated at 4 °C (black), 90 °C (red), and 99 °C (blue) for 20 min before loaded onto an analytical column for gel filtration. The retention profile of RBD was monitored by intrinsic tryptophan fluorescence. (**D**) Three disulfide bonds (orange spheres) segregate the two motifs (α_383-388_ and β_367-370_, green) from the RBM (orange cycle). The two SR31 CDRs are shown as wheat cartoon. α_383-388_ is tethered between Cys379/432 and Cys391/525; β_367-360_ is tethered between Cys379/432 and Cys336/361.

Remarkable, the dramatic rearrangements did not cause noticeable conformational change of RBM (**Fig. 4A**) nor did it affect ACE2 binding (**Fig. 1C**). Given that RBD is a relatively small entity, and that the two surfaces are relatively close (∼25 Å), this was somewhat unexpected. A probable explanation is that RBD is very rigid and hence stable. Indeed, as shown in **Fig. 4C**, RBD showed ultra-stability, with an apparent melting temperature of greater than 95 °C (20-min heating).

Intriguingly, the rearrangement happens at a region that is rich in disulfide bonds. Specifically, β_367-370_ is tethered between the disulfide pairs Cys379-Cys432 and Cys336-Cys361, and α_383-388_ bridges Cys379-Cys432 and Cys-391-Cys525 (**Fig. 4D**). Thus, the three disulfide bonds segregate the two local motifs from the rest of RBD, preventing these conformational changes from propagating through the domain.

### SR31 as a non-competing sybody for RBM binders

The neutral feature of SR31 so far suggests it could bind to RBD in addition to RBM binders such as MR17 and SR4 (*35*). Indeed, BLI assays showed no competition between SR31 and MR17 (**Fig. 5A**), indicating a ‘sandwich complex’ where the RBD is bound with both sybodies. This non-competing feature was also observed in the case of MR6 (**Fig. 5B**) which has also been shown to have neutralizing activities (*35*). As a further proof for the simultaneous binding, we determined the structure of the sandwich complex SR31-RBD-MR17 (**Fig. 5C, Table 1**) to 2.10 Å resolution. The sandwich complex was similar to the individual MR17- and SR31-RBD complexes, with an overall Cα RMSD of 0.667 and 0.375 Å, respectively. Aligning the sandwich complex with the MR17-RBD structure revealed no noticeable changes at the MR17-binding surface (**Fig. 5C**), reinforcing the idea that SR31-binding does not allosterically change the RBM surface nor affect RBM binders.

**Fig. 5.**
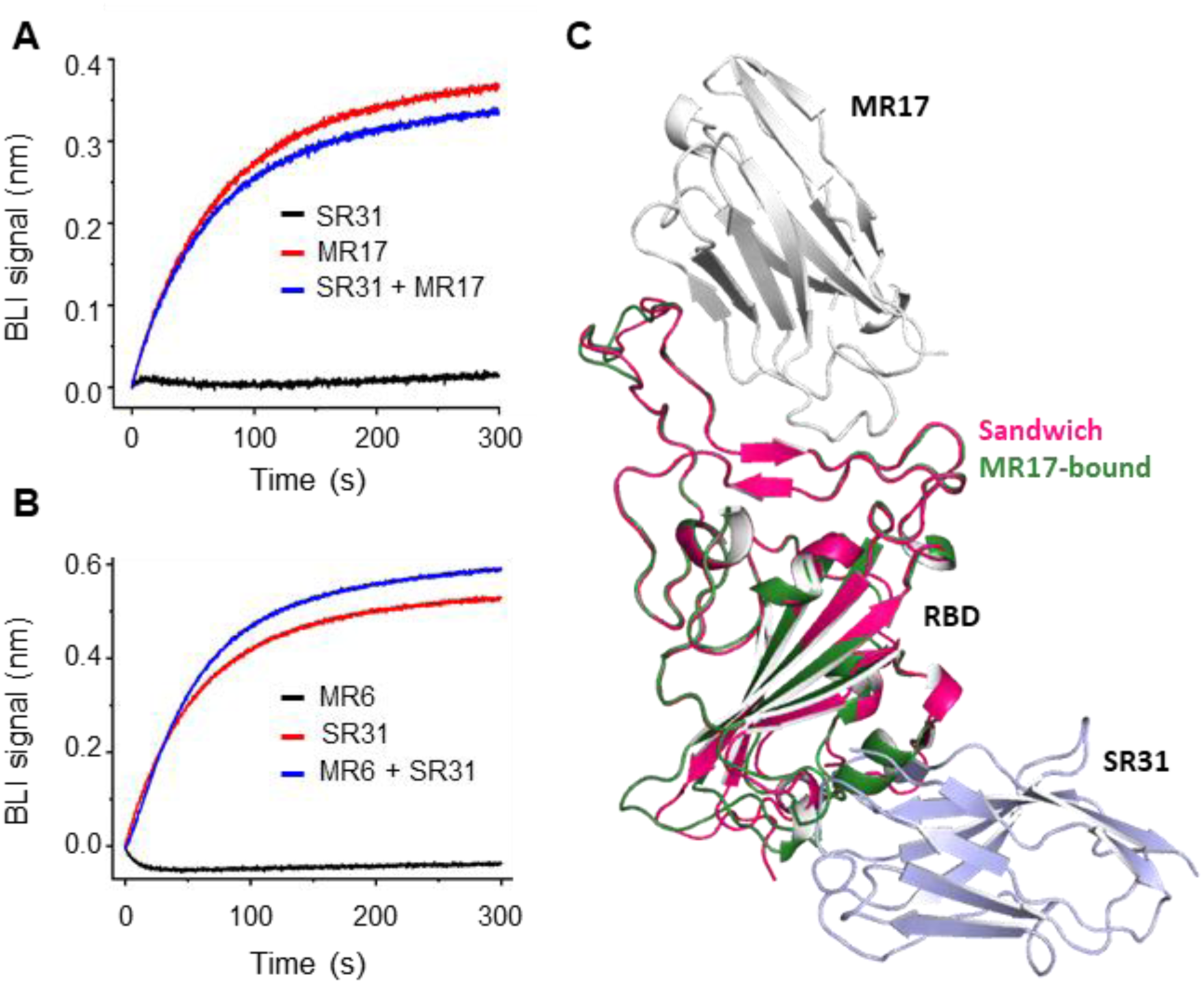
SR31 could pair with RBM nanobodies to bind RBD. (**A, B**) SR31 does not interfere with MR17 (**A**) or MR6 (**B**) for RBD-binding. In **A**, a RBD-coated sensor was pre-saturated in 200 nM of SR31 before incubating with SR31 alone (black) or a mixture (blue) of SR31 and MR17/MR6. In **B**, the sensor was saturated with MR6 before analyzed with SR31. For control purposes, the binding between RBD and the sybody used in the pre-incubation was also characterized (red). (**C**) Alignment of the sandwich complex structure containing MR17 (grey), RBD (red), and SR31 (light blue) to the two-component complex structure (RBD (green) and MR17, PDB ID 7c8w) (*35*).

### SR31 fusion increases affinity and neutralization activity of MR17 and MR6

Although SR31 does not neutralize SARS-CoV-2 pseudovirus itself, its high-affinity may help increase the affinity of other neutralizing nanobodies through avidity effect by fusion. Indeed, the biparatopic fusion SR31-MR17 displayed remarkable increase in binding affinity compared to SR31 or MR17 alone. Its *K*_D_ of 0.3 nM (**Fig. 6A**) was lower than MR17 (*K*_D_ = 83.7 nM) (*35*) by 230 folds and lower than SR31 (*K*_D_ = 5.6 nM) by 17 folds. Consistently, SR31-MR17 neutralized SARS-CoV-2 pseudovirus 13 times more effectively (in molarity) than MR17 alone (**Fig. 6B**).

**Fig. 6.**
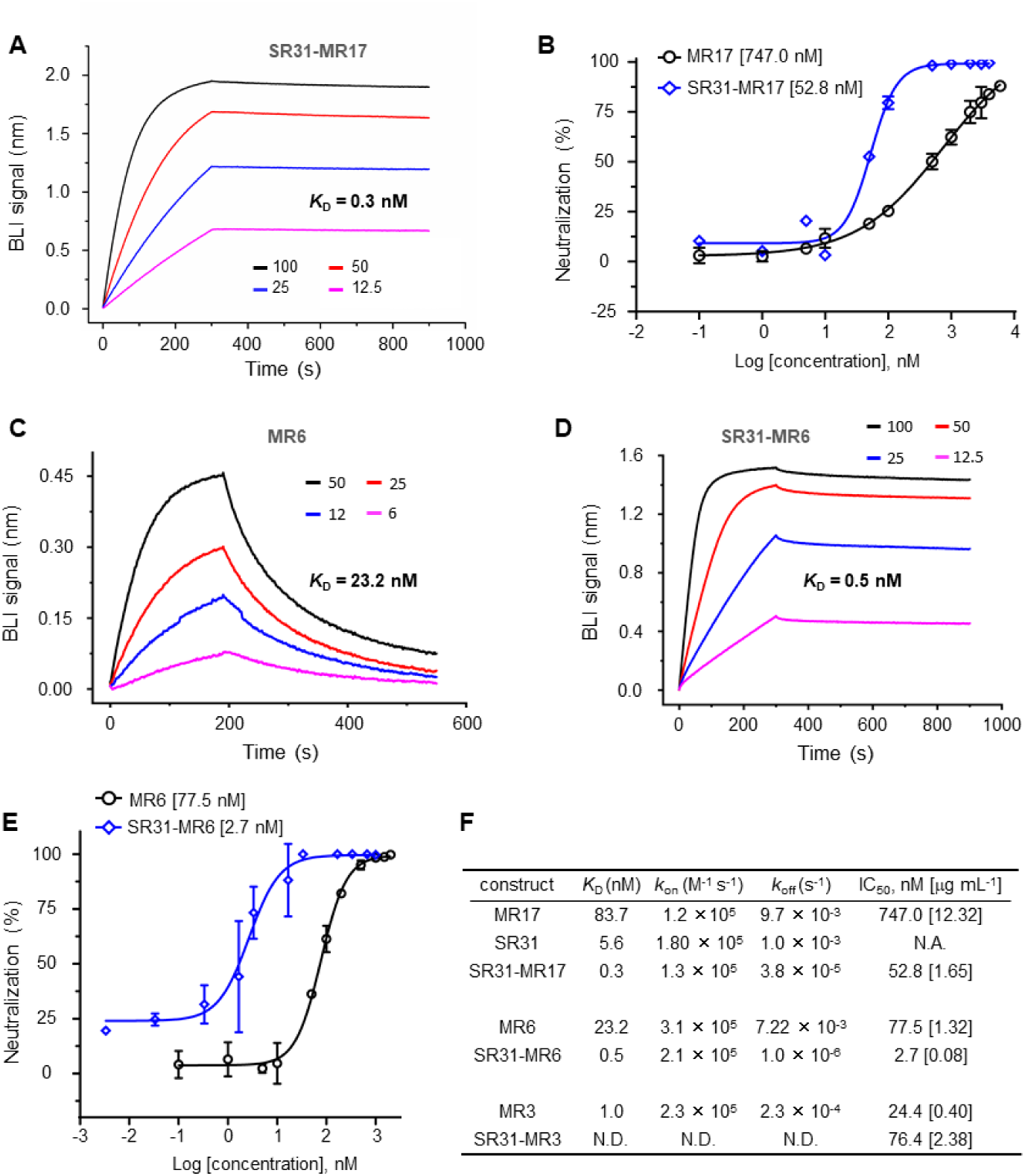
SR31 increases binding affinity and neutralization activity of two fusion partners. (**A**) BLI binding assay with immobilized RBD and the biparatopic sybody SR31-MR17 as analyte at increasing concentrations (nM). (**B**) Neutralization assay of MR17 (black) and SR31-MR17 (blue). (**C, D**) Binding kinetics for the RBD-binding by MR6 (**C**) or by SR31-MR6 (**D**). Concentrations (nM) of the sybodies used in the binding assay are labeled. (**E**) Neutralization assay of MR6 (black) and SR31-MR6 (blue) using SARS-CoV-2 pseudovirus. (**F**) Summary of the comparison between monovalent sybodies and divalent (biparatopic) sybodies for binding kinetics and neutralization activities.

That SR31 can enhance potency of its fusion partner was also demonstrated in the case for MR6. At its free form, MR6 bound to RBD with a *K*_D_ of 23.2 nM (**Fig. 6C**), and showed modest neutralizing activity with an IC_50_ of 1.32 μg mL^-1^ (77.5 nM). Fusing it to SR31 increased its affinity by over 40 folds, displaying a *K*_D_ of 0.5 nM (**Fig. 6D**). In line with this, SR31-MR6 showed a 27-fold higher neutralization activity compared to MR6, with an IC_50_ of 2.7 nM (0.08 μg mL^-1^) (**Fig. 6E**). Interestingly, when fused to MR3, a neutralizing antibody that had higher affinity (*K*_D_ = 1.0 nM) than SR31, the neutralizing activity decreased by 2 folds (**Fig. 6F**). Possible reasons include steric incompatibility caused by improper link length, and allosteric effects. Such hypothesis warrants future structural investigation.

Binding affinity and neutralizing activity are important characteristics of therapeutic antibodies. For modestly neutralizing nanobodies, the potency can be increased in a number of ways, including random mutagenesis (*22*), structure-based design (*35*), and fusion (*35, 36, 42*). Compared with the other two approaches, the fusion technique is more rapid, less involving and does not rely on prior structural information.

Depending on whether the two fusion partners are the same, divalent nanobodies can be categorized into two types: monoparatopic and biparatopic. Biparatopic fusions recognize two distinct epitopes on the same target. Therefore, they are more likely to be resistant to escape mutants because simultaneous mutations at two epitopes should occur at a much lower rate than at a single epitope.

Because of the minute size, SR31 could be used as an ‘add-on’ to monoclonal antibodies, scFv fragments, and other nanobodies to enhance their affinity and potency, especially for those with modest neutralizing activities. In addition, due to its small size and high stability, SR31 may be chemically modified as a vector to deliver small-molecule inhibitors specifically targeting SARS-CoV-2.

In summary, we have structurally characterized SR31, a high-affinity nanobody against SARS-CoV-2 RBD. Although lacking neutralizing activity alone, SR31 is an attractive biparatopic partner for RBM-binders owing to its distinct epitope from RBM. Our work presents a generally useful strategy and offers a simple and fast approach to enhance potency of modestly active antibodies against SARS-CoV2.

## ACKNOWLEDGMENTS

We thank the staff members of the Large-scale Protein Preparation System for equipment maintenance and management, at the SSRF-BL18U1 and BL19U1 beamlines at National Facility for Protein Science (Shanghai) for technical support and assistance. This work has been supported by the Strategic Priority Research Program of CAS (XDB37020204, D.Li), Key Program of CAS Frontier Science (QYZDB-SSW-SMC037, D.Li), CAS Facility-based Open Research Program, the National Natural Science Foundation of China (31870726, D.Li), the National Key R&D Program of China (2020YFC0845900, D.La.), CAS president’s international fellowship initiative (2020VBA0023, D.La.) and Shanghai Municipal Science and Technology Major Project (20431900402, D.La.).

## AUTHOR CONTRIBUTIONS

H.Y., H.C. & T.L. characterized and crystalized the sybody-RBD complexes. B.Z. performed neutralization assays under supervision of D.La.. D.Li & W.Q. collected X-ray diffraction data. D.Li determined and analyzed structures. D.Li wrote the manuscript with inputs from H.Y., H.C., & T.L.

## MATERIALS AND METHODS

### Protein purification

SARS-CoV-2 RBD was expressed essentially as described (*35*). Briefly, a DNA fragment encoding, from N- to C-terminus, residues 330-541 of SARS-CoV2 S, a Gly-Thr linker, the 3C protease site (LEVLFQGP), a Gly-Ser linker, the Avi tag (GLNDIFEAQKIEWHE), a Ser-Gly linker, and a deca-His tag were cloned into the pFastBac-based vector. Baculovirus was generated in *Sf9* cells following the Invitrogen Bac-to-Bac transfection protocol. High Five insect cells were infected with P3 virus. Medium was collected 48-60 h post infection and incubated with 3.0 mL of Ni-Sepharose Excel (Cat 17-3712-03, GE Healthcare) pre-equilibrated with **Buffer A** (150 mM NaCl, 20 mM Tris pH8.0). After batch binding for 2-3 h, the resin was washed with 20 mM of imidazole in **Buffer A** and eluted with 300 mM imidazole in **Buffer A**.

C-terminally His-tagged sybodies were expressed in *Escherichia coli* MC1061 cells. Cells carrying sybody-encoding genes in the vector pSb-init (*41, 43*) were grown in Terrific Broth (TB, 0.17 M KH_2_PO_4_ and 0.72 M K_2_HPO_4_, 1.2 %(w/v) tryptone, 2.4 %(w/v) yeast extract, 0.5% (v/v) glycerol) plus 25 mg L^-1^ chloramphenicol to OD_600_ of 0.5 at 37 °C. Cells were allowed to grow for another 1.5 h at 22 °C before induced with 0.02 %(w/v) arabinose for 17 h. Cells were harvested and lysed by osmotic shock as follows. Cell suspension in 20 mL of TES-high Buffer (0.5 M sucrose, 0.5 mM EDTA, and 0.2 M Tris-HCl pH 8.0) was first incubated at 4 °C for 30 min for dehydration. To the cell suspension, 40 mL of ice-cold MilliQ H_2_O was added for rehydration at 4 °C for 1 h. The suspension was centrifuged at 20,000×g at 4 °C for 30 min to collect supernatant which contained periplasmic extracts. Appropriate buffers were added to the supernatant to have a final concentration of 150 mM NaCl, 2 mM MgCl_2,_ and 20 mM imidazole. The supernatant was then incubated with Ni-NTA resin pre-equilibrated with 20 mM of imidazole in **Buffer B** (150 mM NaCl and 20 mM Tris HCl pH 8.0). After batch-binding for 2 h, the Ni-NTA resin were subsequently washed and eluted with 30 mM and 300 mM imidazole in **Buffer B**, before desalted into PBS buffer.

Divalent sybodies were engineered to have SR31 at the N-terminal and other sybodies at the C-terminal. The DNA fragments were linked together with sequences encoding Gly-Ser linkers at specified length by Gibson Assembly and insertion PCR. Divalent sybodies were expressed and purified essentially as for the monovalent sybodies.

For crystallization, SR31 or SR31-MR17 was mixed with RBD at a 1:1.5molar ratio. The mixture was then loaded onto a Superdex 200 column for gel filtration. Fractions containing the complex were pooled and concentrated to 10 mg mL^-1^.

### Fluorescence-detection size-exclusion chromatography (FSEC)

To screen RBD binders by size exclusion chromatography (SEC) using unpurified sybodies, RBD was fluorescently labelled as follows. First the avi-tagged RBD was enzymatically biotinylated. It was then incubated with fluorescein-labeled streptavidin. The bright fluorescence of the RBD-streptavidin complex at visible wavelength enables convenient and specific monitoring of RBD SEC behavior without the need for purification.

To assess if sybody of interests binds RBD, purified or unpurified sybody was mixed with the fluorescent RBD before injecting on an analytical SEC column connected to a HPLC machine equipped with a fluorescence detector. The retention profile was then recorded by the fluorescence signal at the excitation/emission pair of 482/508 nm.

### Bio-layer interferometry assay

Bio-layer interferometry (BLI) was used to measure binding kinetics between sybodies and RBD. Biotinylated RBD (2 μg mL^-1^ in 0.005 %(v/v) Tween 20, 150 mM NaCl, 20 mM Tris HCl pH 8.0) was first bound to the SA sensor (Cat 18-5019, ForteBio) which was coated with streptavidin. The sensor was equilibrated (baseline) for 120 s at 30 °C. The sensor was then soaked with sybodies at various concentrations (association) for 200-300 s, before moving into sybody-free buffer for dissociation. BLI signal was monitored during the whole process. Data were fitted with a 1:1 stoichiometry using the build-in software Analysis 10.0 for kinetic parameters. For competitive assay of the RBD between SR31 and ACE2, the RBD-coated sensor was saturated in 200 nM of SR31, before soaked in 25 nM SR31 with or without 25 nM of ACE2. As a control, BLI assays were also carried out by soaking the RBD-coated sensor in ACE2 without SR31. For competitive RBD-binding assays for different sybodies, the assays were carried out the same manner as described above.

### Pseudotyped particle production and neutralizing assay

To generate retroviral pseudotyped particles, HEK293T cells were co-transfected with the vectors expressing the various viral envelope glycoproteins, the murine leukemia virus core/packaging components (MLV Gag-Pol), and a retroviral transfect vector expressing the green fluorescence protein (GFP). The S protein of SARS-CoV and SARS-CoV2 in the vector phCMV were truncated by 19 amino acids at the C-terminus. Pseudotyped particles generated this way were harvested 48 h post-transfection by centrifugation and the supernatant was filtered through a 0.45-μm membrane before applying for neutralization assays.

Fifty microliters of VeroE6-hACE2 cells (10^4^ cells/well) were seeded in a 48-well plate. After 24 h, cells were infected with 100 μL of pseudovirus prepared above. When sybodies were included, they were incubated with the pseudovirus for 1 h at 37 °C before infection. The supernatant of cell culture was removed 6 h after infection and replaced with medium. Cells were allowed to grow for 72 h at 37 °C. GFP expression level, as an indication of infectivity, was monitored by fluorescence-activated flow cytometry analysis.

After 6 h of co-incubation, the supernatants were removed and the cells were incubated in medium (Dulbecco’s modified Eagle’s medium-2% fetal calf serum) for 72 h at 37 °C. GFP expression was determined by fluorescence -activated flow cytometry analysis. Cells incubated with medium-only were used as a control to calculate percent inhibition.

### Crystallization

Crystallization experiments were conducted using a Gryphon LCP robot. A two-well sitting-drop plate was filled by 70 μL of precipitant solution as the reservoir. To each well, 100 nL of protein solution was touch-dispensed using the LCP dispenser of the robot. The protein solution was then mixed with 100 nL of precipitant solution delivered by the 96-headed tips. Plates were sealed with transparent tape (Cat HR4-506, Hampton research) and incubated in a Rocker Imager 1000 at 20 °C for automated imaging.

Crystals for the SR31-RBD complex were grown in 2.0 M Sodium formate, 0.1 M Sodium acetate trihydrate pH 4.6. Cryo protection was achieved by adding 20 %(v/v) glycerol to the mother liquor condition. Crystals for the SR31-MR17-RBD complex were grown in 0.1 M cadmium chloride, 0.1 M Na-acetate pH 4.5, 30 % PEG 400, 4% v/v (±) -1,3-butanediol. Cryo protection was achieved by adding 20 %(v/v) glycerol in the mother liquor condition.

Desired crystals were cryo-protected, harvested using a MiTeGen loop under a microscope, and flash-cooled in liquid nitrogen before diffraction.

### Data collection and structure determination

X-ray diffraction data were collected at beamline BL19U1 (*44*) at Shanghai Synchrotron Radiation Facility with a 50 × 50 μm beam on a Pilatus 6M detector, with oscillation of 0.5° and a wavelength of 0.97853 Å. Data were integrated using the software XDS (*45*), and scaled and merged using Aimless (*46*). The SR31-RBD structure was solved by molecular replacement using Phaser (*47*) with PDB IDs 6M0J and 5M13 (*41*) as the search model. The SR31-MR17-RBD structure was solved using the SR31-RBD and MR17 structure (*35*) as search models. The models were manually adjusted as guided by the 2F_o_-F_c_ maps in Coot (*48*), and refined using Phenix (*49*). Structures were visualized using PyMol (*50*).

## Data availability

The structure factors and coordinates were deposited in the protein data bank (PDB) under accession codes 7D2Z (SR31+RBD) and 7D30 (SR31-MR17+RBD).

---

**Table S1.**
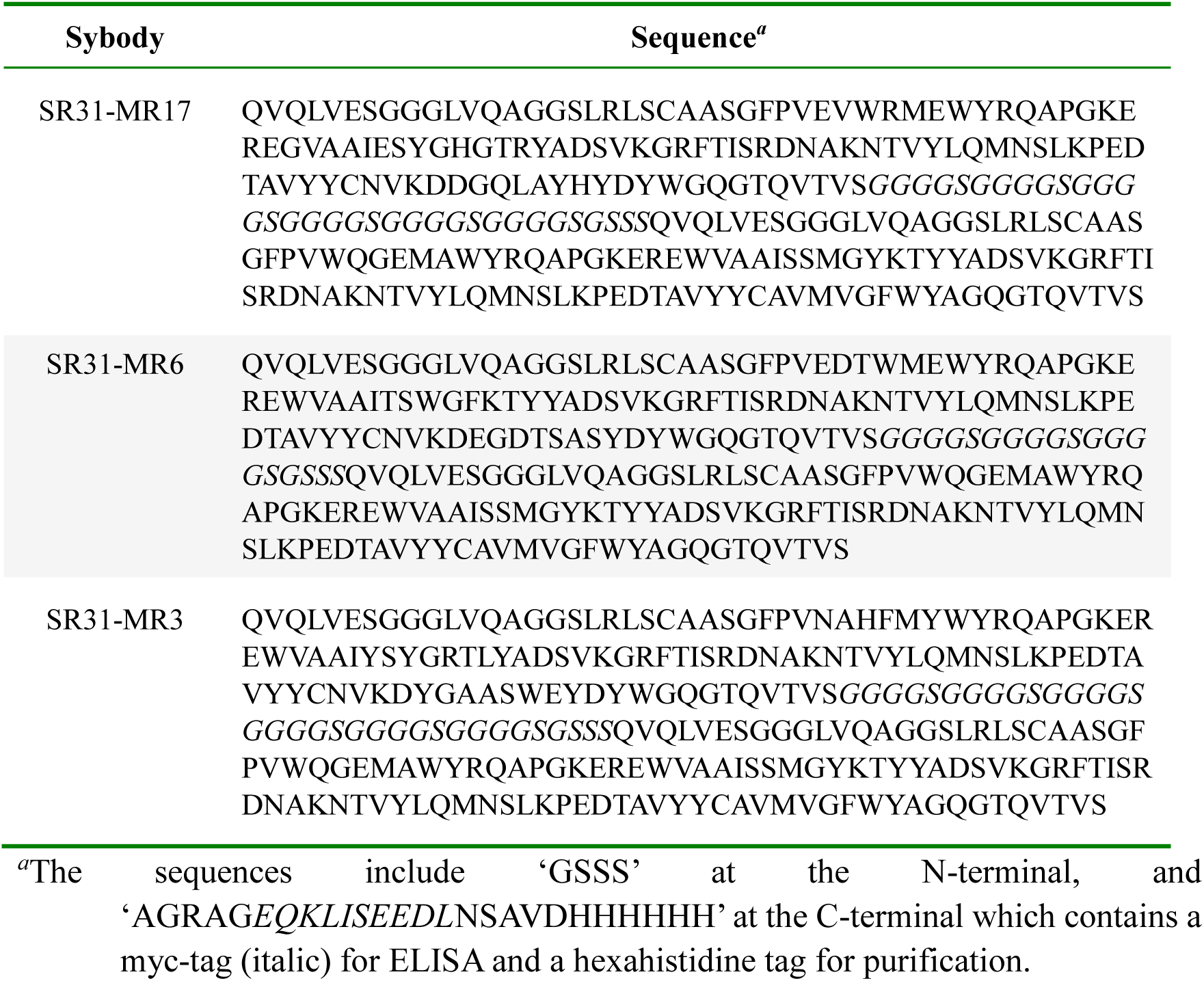
Sequences of biparatopic sybodies.

